# A DEAD BOX RNA helicase from *Medicago truncatula* is hijacked by an RNA-binding effector from the root pathogen *Aphanomyces euteiches* to facilitate host infection

**DOI:** 10.1101/2020.06.17.157404

**Authors:** L. Camborde, A. Kiselev, M.J.C. Pel, A. Leru, A. Jauneau, C. Pouzet, B. Dumas, E. Gaulin

## Abstract

Microbial effectors from plant pathogens are molecules that target host components to facilitate colonization. While eukaryotic pathogens are virtually able to produce hundreds of effectors, the molecular mechanisms allowing effectors to promote infection are still largely unexplored. Here we show that the effector AeSSP1256 from the soilborne oomycete pathogen *Aphanomyces euteiches* is able to interact with plant RNA. Heterologous expression of AeSSP1256 delays *Medicago truncatula* host roots development and facilitate pathogen colonization. Transcriptomic analyses of AeSSP1256-expressing roots show a downregulation of genes implicated in ribosome biogenesis pathway. A yeast-two hybrid approach reveals that AeSSP1256 associates with a nucleolar L7 ribosomal protein and a *M. truncatula* RNA helicase (MtRH10) orthologous to the Arabidopsis RNA helicase RH10. Association of AeSSP1256 with MtRH10 impaired the capacity of MtRH10 to bind nucleic acids. Promoter:GUS composite plants revealed that MtRH10 is expressed preferentially in the meristematic root cells. Missense MtRH10 plants displayed shorter roots with developmental delay and are more susceptible to *A. euteiches* infection. These results show that the effector AeSSP1256 facilitates pathogen infection by causing stress on plant ribosome biogenesis and by hijacking a host RNA helicase involved in root development and resistance to root pathogens.

## Introduction

Plant pathogens divert host cellular physiology to promote their own proliferation by producing effector proteins that interact with molecular targets (Gaulin et al., 2018). Numerous studies indicate large variation in the effector repertoire of plant pathogens suggesting that a large number of molecular mechanisms are targeted.

Oomycetes constitute a large phylum that includes important eukaryotic pathogens, and many of which are destructive plant or animal pathogens (Kamoun et al., 2015; van West and Beakes, 2014). They share common morphological characteristics with true fungi as filamentous growth, osmotrophic feeding or the presence of a cell wall, but they evolved independently (Judelson, 2017). Oomycetes are included in the Stramenopile lineage and have diatoms and brown algae as closest cousins. These filamentous microorganisms have the capacity to adapt to different environment as illustrated by their capacity to develop resistance to anti-oomycete chemicals or quickly overcome plant resistance (Rodenburg et al., 2020).

Comprehensive identification of oomycete proteins that act as effectors is challenging. Up to now, computational predictions of effector proteins have provide a fast approach to identify putative candidate effectors in oomycetes (Haas et al., 2009; Tabima and Grünwald, 2019). Based on their predictive subcellular localization within the host cells they are classified as extracellular (apoplasmic) or intracellular (cytoplasmic) effectors. As example, RxLR and Crinklers (CRNs) constitute the two largest family of oomycetes intracellular effectors that contain hundreds of members per family (McGowan and Fitzpatrick, 2017). While oomycete effector proteins have probably different mechanisms of action, what they have in common might be the ability to facilitate pathogen development. Nonetheless, computational predictions do not give any clues regarding the putative role of theses effectors since numerous effectors are devoid of any functional domains. Therefore, biochemical and molecular studies are used to discover and confirm the functional activity of these proteins. To promote infection oomycete intracellular effectors interfere with many host routes which include for example signaling such as MAPKinase cascades (King et al., 2014), phytohormone-mediated immunity (Boevink et al. 2016; Liu et al. 2014), trafficking vesicles secretion (Du et al., 2015) or autophagosome formation (Dagdas et al., 2016). Growing evidences point to plant nucleus as an important compartment within these interactions thanks to the large portfolio of putative nucleus-targeted effectors predicted in oomycete genomes. The study of subcellular localization of fifty-two *Phytophthora infestans* RxLR effectors upregulated during the early stage of host infection show that nucleocytoplasmic distribution is the most common pattern, with 25% effectors that display a strong nuclear association (Wang et al. 2019). The CRN family was firstly reported as a class of nuclear effector from *P. infestans* (Schornack et al., 2010), around 50% of predicted NLS-containing CRN effectors from *P. capsici* showed nuclear localization (Stam et al., 2013) and numerous CRNs effectors from *P. sojae* such as PsCRN108, PsCRN63 or PsCRN115 harbor a nuclear localization (Song et al., 2015; Zhang et al., 2015). In agreement with this, different mechanisms of action at the nuclear level have been reported for oomycete effectors such as the alteration of genes transcription (Wirthmueller et al., 2018; Song et al., 2015; He et al., 2019), the mislocalisation of transcription factor (Mclellan et al., 2013), the suppression of RNA silencing by inhibition of siRNA accumulation (Qiao et al., 2015; Xiong et al., 2014) or the induction of plant DNA-damage (Camborde et al. 2019; Ramirez-Garcés et al. 2016). However specific function has been assigned to very few effectors.

We previously use comparative genomics and predictive approaches on the *Aphanomyces* genus to identify putative effectors and characterized a large family of small secreted proteins (SSPs) (Gaulin et al., 2018). SSPs harbor a predicted N secretion signal, are less than 300 residues in size and devoid of any functional annotation. More than 290 SSPs are predicted in the legume pathogen *A. euteiches* (AeSSP) while 138 members with no obvious similarity to AeSSP members are reported in the crustacean parasite *A. astaci* (Gaulin et al., 2018). This specific SSP repertoire suggests its role in adaption of *Aphanomyces* species to divergent hosts. We have previously identified one AeSSP (AeSSP1256) based on a screen aiming to identify SSP able to promote infection of *Nicotiana benthamiana* plants by the leaf pathogen *Phytophthora capsici*. AeSSP1256 harbors a nuclear localization signal indicating its putative translocation to host nucleus. However, the function of this protein remained to be identified.

Here we report on the functional analysis of AeSSP1256 and the characterization of its plant molecular target. We show that AeSSP1256 binds RNA *in planta*, induces developmental defects when expressed in *M. truncatula* roots and promotes *A. euteiches* infection. This phenotype is correlated with a downregulation of a set of ribosomal protein genes. A yeast two-hybrid approach identified a host RNA helicase (MtRH10) and a L7 ribosomal protein as interactors of AeSSP1256. By FRET-FLIM analyses we reveal that AeSSP1256 co-opts MtRH10 to abolish its nucleic acid binding capacity. We provide a mechanistic explanation of this observation by demonstrating the implication of MtRH10 in roots development by generating missense and overexpressing *Medicago* lines. Finally we observed that silenced-MtRH10 roots are highly susceptible to *A. euteiches* infection like AeSSP1256-expressing roots, showing that MtRH10 as AeSSP1256 activities modify the outcome of the infection. We now present results supporting effector-mediated manipulation of a nuclear RNA helicase as a virulence mechanism during plant-eukaryotic pathogens interactions.

## Results

### AeSSP1256 contains RGG/RG domains and binds RNA *in planta*

AeSSP1256 is a member of a large family of *A. euteiches* effectors devoid of any predicted functional domain, except the presence of a signal peptide at the N-terminus (Gaulin et al., 2018). As showed in **Figure 1A**, AeSSP1256 protein is enriched in glycine residues (30% of the amino acid sequence). Analysis using the Eukaryotic Linear Motif database (Gouw et al., 2018) revealed 3 GGRGG motifs (positions 81-85; 95-99 and 99-103). These motifs are variant arginine methylation site from arginine-glycine(-glycine) (RGG/RG) domains, presents in many ribonucleoproteins and involved in RNA binding (Thandapani et al., 2013; Bourgeois et al., 2020). We then noticed the presence of two di-RGG domains (RGG(X_0-5_)RGG) (position 75-85 and 97-103) and one di-RG domains (RG(X_0-5_)RG) (position 123-126) corresponding to RGG or RG motifs that are spaced less than 5 residues (Chong et al., 2018). According to RGG/RG definition, those repeats occur in low-complexity region of the protein (position 60-180) (Chong et al., 2018) and are associated with di-glycine motifs and GR or GGR sequences (**Figure 1A**), which are also common in RGG/RG-containing proteins (Chong et al., 2018). Considering that RGG/RG domains are conserved from yeast to humans (Rajyaguru and Parker, 2012) and represent the second most common RNA binding domain in the human genome (Ozdilek et al., 2017), we therefore investigated the RNA binding ability of AeSSP1256. To test this, we performed a FRET-FLIM assay on *N. benthamiana* agroinfiltrated leaves with AeSSP1256:GFP fusion protein in presence or absence of Sytox Orange to check its capacity to bind nucleic acids (Camborde et al. 2017). Briefly AeSSP1256:GFP construct is transiently express in *N. benthamiana* leaves where it accumulates in the nucleus (Gaulin et al., 2018). Samples are collected 24h after treatment and nucleic acids labeled with the Sytox Orange dye. In presence of Sytox, if the GFP fusion protein is in close proximity (<10nm) with nucleic acids, the GFP lifetime of the GFP tagged protein will significantly decrease, due to energy transfer between the donor (GFP) and the acceptor (Sytox). To distinguish RNA interactions from DNA interactions, an RNase treatment can be performed. In the case of a specific RNA-protein interaction, no FRET acceptor will be available due to RNA degradation and the lifetime of the GFP tagged protein will then return at basal values. It appeared that GFP lifetime of AeSSP1256:GFP decreased significantly in presence of Sytox Orange as reported in ***table 1*** and in **Figure 1B**, decreasing from 2.06 +/- 0.02 ns to 1.84 +/- 0.03 ns. This indicates that AeSSP1256 is able to bind nucleic acids. After an RNase treatment, no significant difference on GFP lifetime was observed in absence (2.01 ns +/- 0.02) or in presence (1.96 ns +/- 0.02) of Sytox Orange, meaning that the FRET was not due to DNA interaction but was specific to RNA (***table 1*** and **Figure 1B**). These results indicate that AeSSP1256 is able to bind nuclear RNA in plant cells.

**Figure 1:**
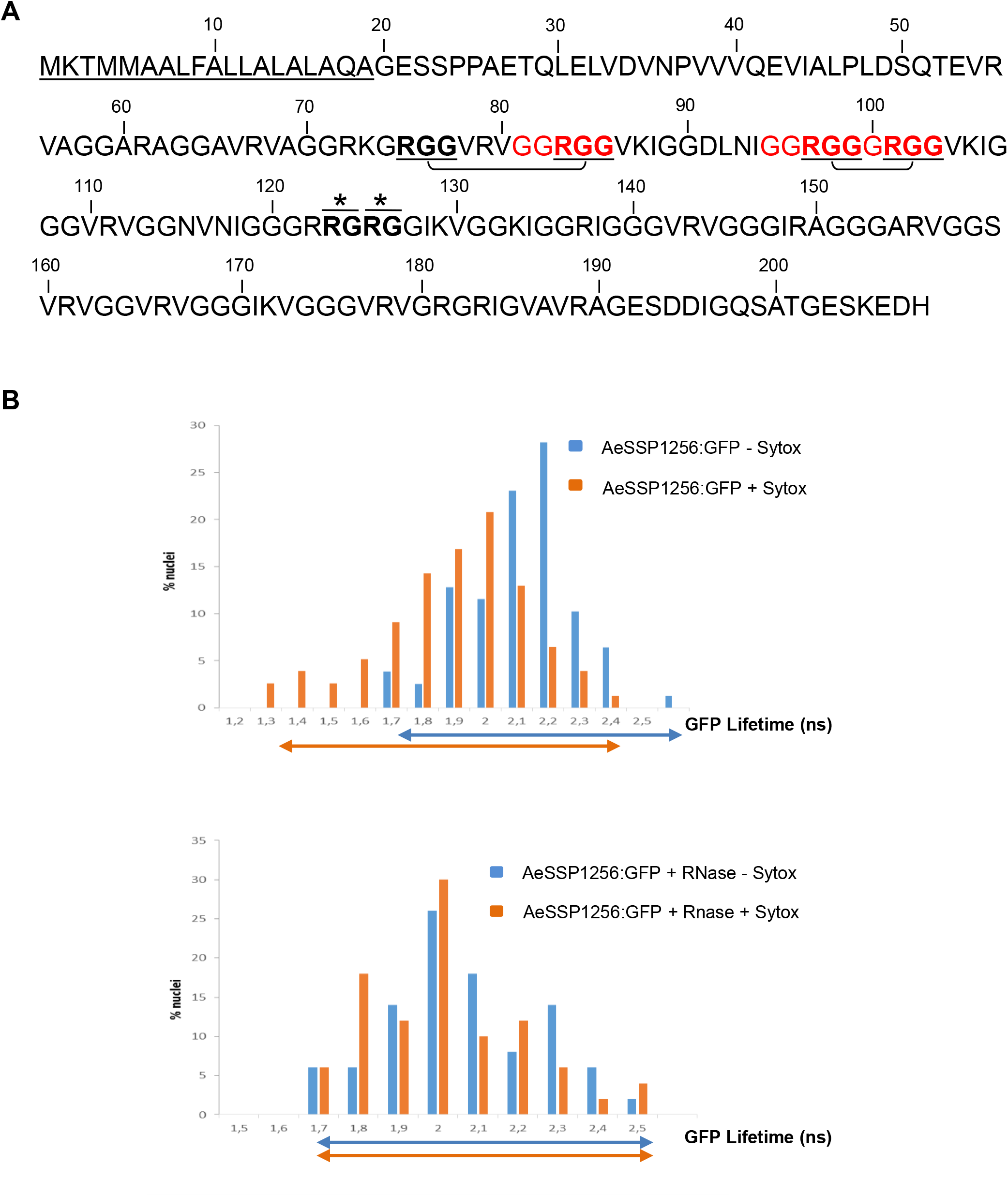
AeSSP1256 is a RNA-binding protein. **(A)** AeSSP1256 protein sequence that shows the signal peptide (underlined), GGRGG boxes (red), RGG domains (bolt, underlined and linked), RG motifs (bolts with asterisks) predicted with Eukaryotic Linear Motif Prediction (Gouw *et al*., 2018). **(B)** One day after agroinfection of *N. benthamiana* leaves with a AeSSP1256:GFP construct, infiltrated area are collected for FRET-FLIM analysis to detect protein/nucleic acid interactions as described by Camborde *et al*., 2018. Without RNAse treatment and in presence of nucleic acids dye Sytox Orange, the AeSSP1256:GFP lifetime decreases to shorter values, indicating that the proteins bounded to nucleic acids (top panel). After RNase treatment, no significant decrease in the GFP lifetime was observed in presence of Sytox Orange, indicating that AeSSP1256:GFP proteins were bounded specifically to RNA (bottom panel). Histograms show the distribution of nuclei (%) according to classes of AeSSP1256:GFP lifetime in the absence (blue bars) or presence (orange bars) of the nucleic acids dye Sytox Orange. Arrows represent GFP lifetime distribution range.

**Table 1:** FRET-FLIM measurements for AeSSP1256:GFP with or without Sytox Orange

| Donor | Acceptor | $\tau$ <sup>(a)</sup> | sem <sup>(b)</sup> | N <sup>(c)</sup> | E <sup>(d)</sup> | <sup>(e)</sup> p-value |
| --- | --- | --- | --- | --- | --- | --- |
| AeSSP1256:GFP | - | 2.06 | 0.020 | 78 | - | - |
| AeSSP1256:GFP | Sytox | 1.84 | 0.026 | 77 | 11 | 1.34E<sup>-09</sup> |
| AeSSP1256:GFP | -<br>(+ RNase) | 2.01 | 0.026 | 50 | - | - |
| AeSSP1256:GFP | Sytox<br>(+ RNase) | 1.96 | 0.027 | 50 | 2.6 | 0.17 |

$\tau$ : mean life-time in nanoseconds (ns). <sup>(b)</sup> s.e.m.: standard error of the mean. <sup>(c)</sup> N: total number of measured nuclei. <sup>(d)</sup> E: FRET efficiency in %: $E=1-(\tau_{DA}/\tau_D)$ . <sup>(e)</sup> p-value (Student's *t* test) of the difference between the donor lifetimes in the presence or absence of acceptor.

### AeSSP1256 impairs *M. truncatula* root development and susceptibility to *A. euteiches*

To check whether expression of AeSSP1256 may have an effect on the host plant, we transformed *M. truncatula* (*Mt*) roots, with a native version of GFP tagged AeSSP1256. As previously observed (Gaulin et al., 2018), confocal analyses confirmed the nuclear localization of the protein in root cells, with accumulation around the nucleolus as a perinucleolar ring (**Figure 2A**) despite the presence of a signal peptide (Gaulin et al., 2018). Anti-GFP western blot analysis on total proteins extracted from transformed roots confirmed the presence of GFP-tagged AeSSP1256 (46.7 kDa expected sizes) (**Figure 2B**). We noticed the presence of a second band around 28 kDa, which is probably free GFP due to the cleavage of the tagged protein. AeSSP1256:GFP transformed plants showed delayed development (**Figure 2C**), with total number of roots and primary root length per plant being significantly lower than values obtained with GFP control plants (**Figure 2D**). As previously observed in *N. benthamiana*, when a KDEL-endoplasmic reticulum (ER) retention signal is added to the native AeSSP1256 construct (Gaulin et al., 2018), AeSSP1256:KDEL:GFP proteins mainly accumulates in the ER (**Supplemental Figure 1A-C**) and roots showed no significant differences in development as compared to GFP control roots (**Supplemental Figure 1D and E**). In contrast a construct devoid of a native signal peptide (SP) shows that the proteins accumulated in root cell nuclei (**Supplemental Figure 1B**), leading to abnormal root development, with symptoms similar to those observed in presence of the AeSSP1256:GFP construct, including shorter primary root and lower number of roots (**Supplemental Figure 1D and E**). Altogether these data show that within the host, AeSSP1256 triggers roots developmental defects thanks to its nuclear localization.

**Figure 2:**
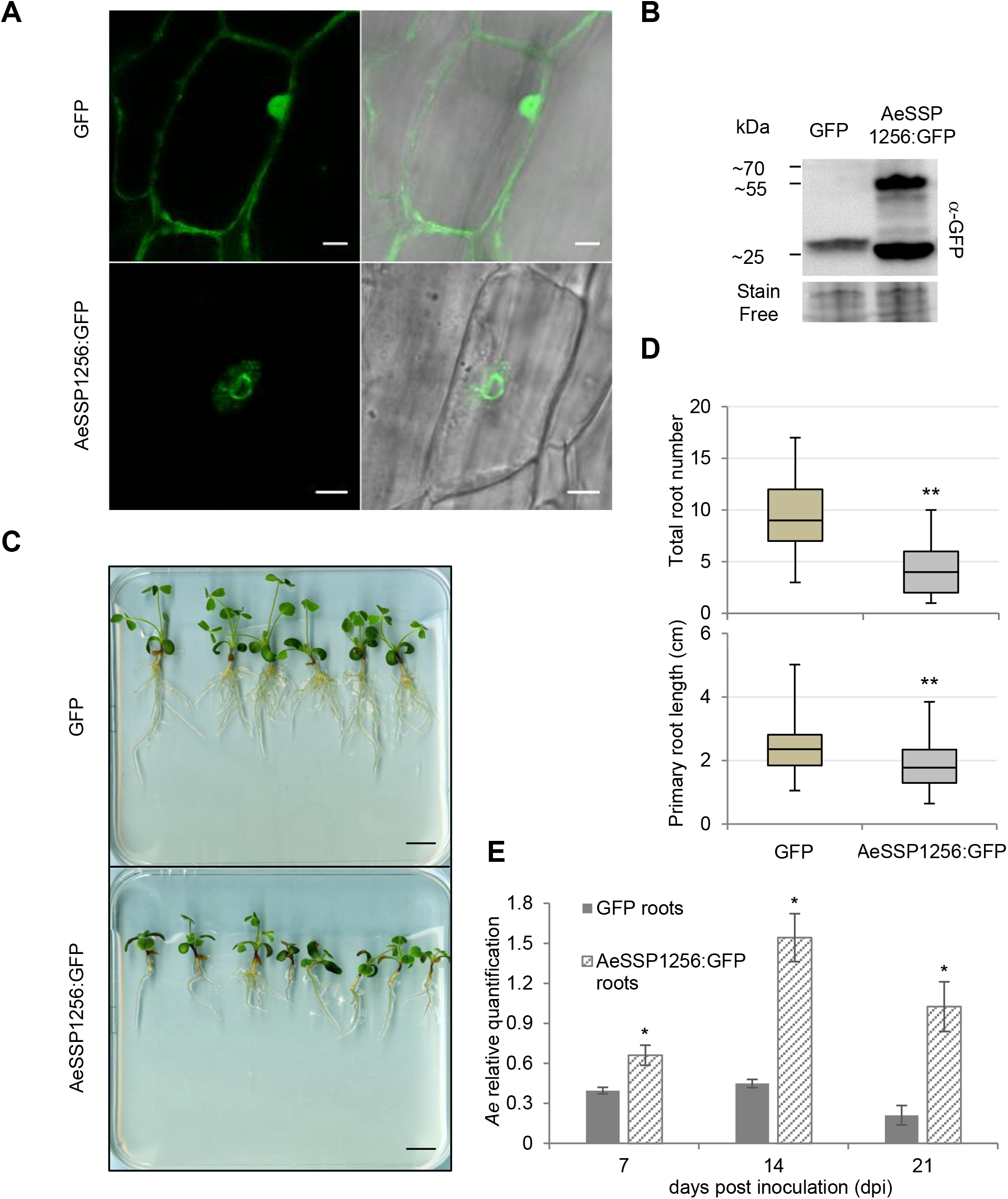
AeSSP1256 pertubs *M. truncatula* roots development and enhances *A. euteiches* susceptibility. *M. truncatula* tolerant A17 lines were transformed using *Agrobacterium rhizogenes-mediated* transformation system to produce GFP or AeSSP1256:GFP composite plants. **(A)** Confocal analysis of *M. truncatula* transformed roots at 21 days after transformation (d.a.t). The GFP control protein presents a nucleocytoplasmic localisation (upper panel), while the AeSSP1256 effector is localized as a ring around the nucleolus (bottom panel). Scale bars: 10μm. **(B)** Total proteins were extracted from transformed *M. truncatula* roots at 21 d.a.t and subjected to western-blot analysis using anti-GFP antibodies. A representative blot shows a band around 28kDa that represents the GFP protein and a band corresponding to the AeSSP1256:GFP protein (expected size 46.5 kDa). **(C)** Representative photographs of AeSSP1256:GFP plants and GFP control plants at 21 d.a.t. Note the reduction in the growth of roots expressing the AeSSP1256 effector as compared to GFP control plants. Scale bar: 1cm. **(D)** Diagram depicting the total root number per plant (upper panel) and primary root length (in cm) per plant (bottom panel) of transformed *M. truncatula* plants at 21 d.a.t. n= 126 plants for GFP and n=79 plants for AeSSP1256:GFP. **(E)** qPCR results showing relative quantification of the *A. euteiches* tubulin gene in *M. truncatula* GFP or AeSSP1256:GFP infected roots at 7, 14 and 21 days post inoculation (d.p.i). For each time point, 45 to 75 plants per construct were used. Asterisks indicate significant differences (Student’s t-test; *: P < 0.05; **: P<0.001).

To investigate whether AeSSP1256 modifies the outcome of the infection, AeSSP1256-transformed roots were inoculated with *A. euteiches* zoospores. RT-qPCR analyses at 7, 14 and 21 days post inoculation were performed to follow pathogen development. At each time of the kinetic, *A. euteiches* is more abundant in *M. truncatula* roots expressing the effector than in GFP control roots (respectively 1.5, 3 and 5 times more) (**Figure 1E**). This indicates that roots are more susceptible to *A. euteiches* in presence of AeSSP1256. Transversal sections of A17-transformed roots followed by Wheat-Germ-Agglutinin (WGA) staining to detect the presence of *A. euteiches*, showed that the pathogen is still restricted to the root cortex either in the presence or absence of AeSSP1256 (**Supplemental Figure 2**). This phenotype is similar to the one observed in the natural A17 *M. truncatula* tolerant line infected by *A. euteiches* (Djébali et al., 2009). This data suggests that defence mechanisms like protection of the central cylinder (Djébali et al., 2009) are still active in AeSSP1256-expressing roots.

### AeSSP1256 affects the expression of genes related to ribosome biogenesis

To understand how AeSSP1256 affects *M. truncatula* roots development and facilitates *A. euteiches* infection, we performed expression analyses by RNASeq using AeSSP1256-expressing roots and GFP controls roots. 4391 genes were differentially express (DE) between the two conditions (*p* adjusted-value <10^−5^) (**Supplemental Table 1a**). Enrichment analysis of ‘Biological process’ GO-terms showed the presence of ‘ribosome biogenesis’ and ‘organonitrogen compound biosynthetic, cellular amide metabolic’ processes terms among the most enriched in AeSSP1256 roots as compared to GFP-expressing roots (**Supplemental Table 1b)**. We noticed that over 90% of DE-genes from ‘ribosome biogenesis’ and ‘translation’ categories are downregulated in AeSSP1256-expressing roots, suggesting that expression of the effector within the roots affects ribosome biogenesis pathway (**Supplemental Table 1a**). To evaluate whether expression of AeSSP1256 mimics infection of *M. truncatula* by *A. euteiches* infection through downregulation of genes related to ribosome biogenesis, we analyzed RNASeq data previously generated on the susceptible F83005.5 *M. truncatula* line nine days after root infection (Gaulin et al., 2018). As shown on the Venn diagram depicting the *M. truncatula* downregulated genes in the different conditions (**Figure 3A, Supplemental Table1c**), among the 270 common downregulated genes between AeSSP1256-expressing roots and susceptible F83-infected lines, 58 genes (>20%) are categorized in the ‘ribosome biogenesis’ and ‘translation’ GO term (**Figure 3B**). We next selected seventeen *M. truncatula* genes to confirm the effect via qRT-PCR. First, we selected ten *A. thaliana* genes related to plant developmental control (i,e mutants with shorter roots phenotype) (**Supplemental Table 1d**) by Blast searches (>80% identity) in A17 line r5.0 genome portal (Pecrix et al., 2018). In addition, seven nucleolar genes coding for ribosomal and ribonucleotides proteins and related to the ‘ribosome biogenesis’ in *M. truncatula* were selected for expression analysis based on KEGG pathway map (https://www.genome.jp/kegg-bin/show_pathway?ko03008) (**Supplemental Table 1d**). As shown on **Figure 3C**, all of the selected genes from *M. truncatula* are downregulated in presence of AeSSP1256, supporting the RNAseq data. Altogether these expression data show that the effector by itself mimics some effects induced by pathogen infection of the susceptible F83 line. At this stage of the study, results point to a perturbation of the ribosome biogenesis pathway of the host plant by the AeSSP1256 effector.

**Figure 3:**
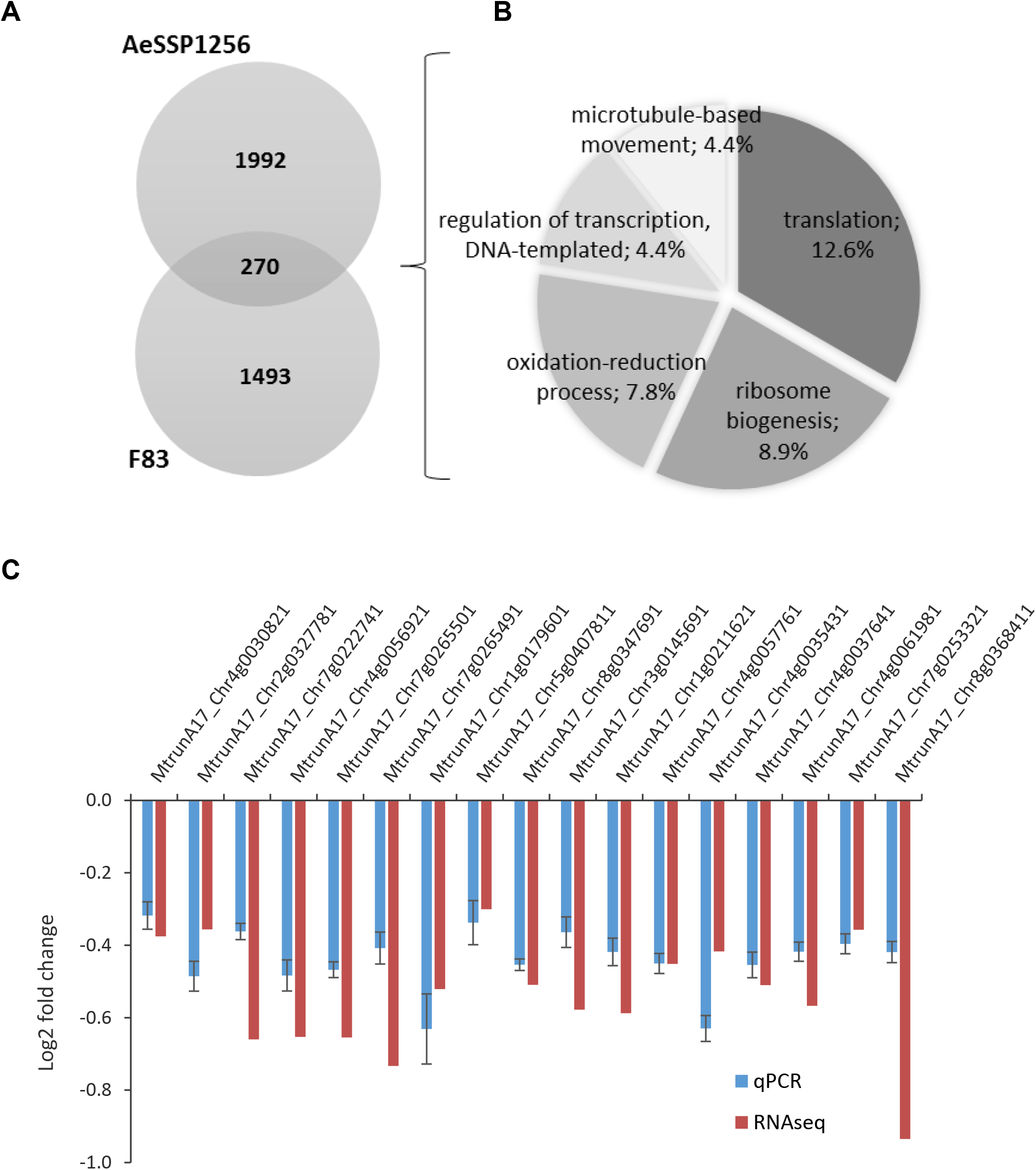
Transcriptomic analyses reveal downregulation of genes related to ribosome biogenesis in both AeSSP1256-expressing roots or *A. euteiches*-infected roots. **(A)** Venn diagram on downregulated genes (number of genes) of two RNASeq experiments: F83 (*M. truncatula* susceptible F83005.5 roots infected by *A.euteiches* at 9 dpi), AeSSP1256 (*M. truncatula* tolerant A17 line expressing AeSSP1256:GFP). **(B)** The most represented GO-terms common between F83-infected line and AeSSP1256-expressing roots of downregulated genes are related to ‘translation and ribosome-biogenesis’. Only GO terms containing more than 10 genes are represented on the pie chart. Numbers on the graph indicate percent of genes with a GO term. **(C)** Comparison of RNASeq (n=4) and qRT-PCR (n=5) on selected ribosome biogenesis-related genes.

### AeSSP1256 targets a DEAD-box RNA helicase and a L7 ribosomal protein

To decipher how AeSSP1256 can affect ribosome biogenesis pathway of the host plant and knowing that numerous RNA-binding proteins interact with protein partners, we searched for AeSSP1256 host protein targets. For this, a Yeast two hybrid (Y2H) library composed of cDNA from *M. truncatula* roots infected with *A. euteiches* was screened with the mature form of the effector. Eight *M. truncatula* coding genes were identified as potential protein targets (**Supplemental Table 2a**), all these genes but one (a lecithin retinol acyltransferase gene) correspond to putative nuclear proteins in accordance with the observed subcellular localization of AeSSP1256.

To confirm the Y2H results, we first expressed AeSSP1256 and candidates in *N. benthamiana* cells to observe their subcellular localization and performed FRET-FLIM experiments to validate protein-protein interactions. Only two candidates showed co-localization with AeSSP1256, a L7 ribosomal protein (RPL7, MtrunA17_Chr4g0002321) and a predicted RNA helicase (RH) (MtrunA17_Chr5g0429221). CFP-tagged version of RPL7 displays a nucleolar localization, with partial co-localization areas in presence of AeSSP1256 (**Supplemental Figure 3A, Table 2b**). FRET-FLIM measurements confirmed the interaction of RPL7:CFP protein with AeSSP1256:YFP effector (**Supplemental Figure 3B, Table 2b**), with a mean CFP lifetime of 2.83 ns +/- 0.03 in absence of the SSP protein, leading to 2.46 ns +/- 0.03 in presence of AeSSP1256:YFP (**Supplemental Table 2b)**.

**Table 2:** FRET-FLIM measurements of CFP:MtRH10 in presence or absence of AeSSP1256:YFP

| Donor | Acceptor | $\tau^{(a)}$ | sem<sup>(b)</sup> | N<sup>(c)</sup> | E<sup>(d)</sup> | <sup>(e)</sup> p-value |
| --- | --- | --- | --- | --- | --- | --- |
| CFP:MtRH10 | - | 2.86 | 0.023 | 50 | - | - |
| CFP:MtRH10 | AeSSP1256:YFP | 2.53 | 0.031 | 31 | 11.1 | 2.56E<sup>-12</sup> |

$\tau$ : mean life-time in nanoseconds (ns). <sup>(b)</sup> s.e.m.: standard error of the mean. <sup>(c)</sup> N: total number of measured nuclei. <sup>(d)</sup> E: FRET efficiency in % : $E=1-(\tau_{DA}/\tau_D)$ . <sup>(e)</sup> p-value (Student's *t* test) of the difference between the donor lifetimes in the presence or absence of acceptor.

The second candidate is a predicted DEAD-box ATP-dependent RNA helicase (MtrunA17_Chr5g0429221), related to the human DDX47 RNA helicase and the RRP3 RH in yeast. Blast analysis revealed that the closest plant orthologs were AtRH10 in *Arabidopsis thaliana* and OsRH10 in *Oryza sativa*. Consequently the *M. truncatula* protein target of AeSSP1256 was named MtRH10. The conserved domains of DEAD-box RNA helicase are depicted in the alignment of MtRH10 with DDX47, RRP3, AtRH10, OsRH10 proteins (**Supplemental Figure 4A**) (Schütz et al., 2010; Gilman et al., 2017). MtRH10 CFP-tagged fusion protein harbors nucleocytoplasmic localization when transiently express in *N. benthamiana* cells (**Figure 4A**), in accordance with the presence of both putative nuclear export signals (NESs) (position 7-37; 87-103; 261-271) and nuclear localization signal (NLS) sequences (position 384-416). When MtRH10 is co-expressed with YFP-tagged version of AeSSP1256, the fluorescence is mainly detected as a ring around the nucleolus, indicating a partial relocalisation of MtRH10 to the AeSSP1256 sites (**Figure 4A**). FRET-FLIM measurements on these nuclei confirm the interaction between AeSSP1256 and the *Medicago* RNA helicase (**Figure 4B)**, with a mean CFP lifetime of 2.86 ns +/- 0.02 in absence of the effector protein, to 2.53 ns +/- 0.03 in presence of AeSSP1256:YFP (**Table 2**).

**Figure 4:**
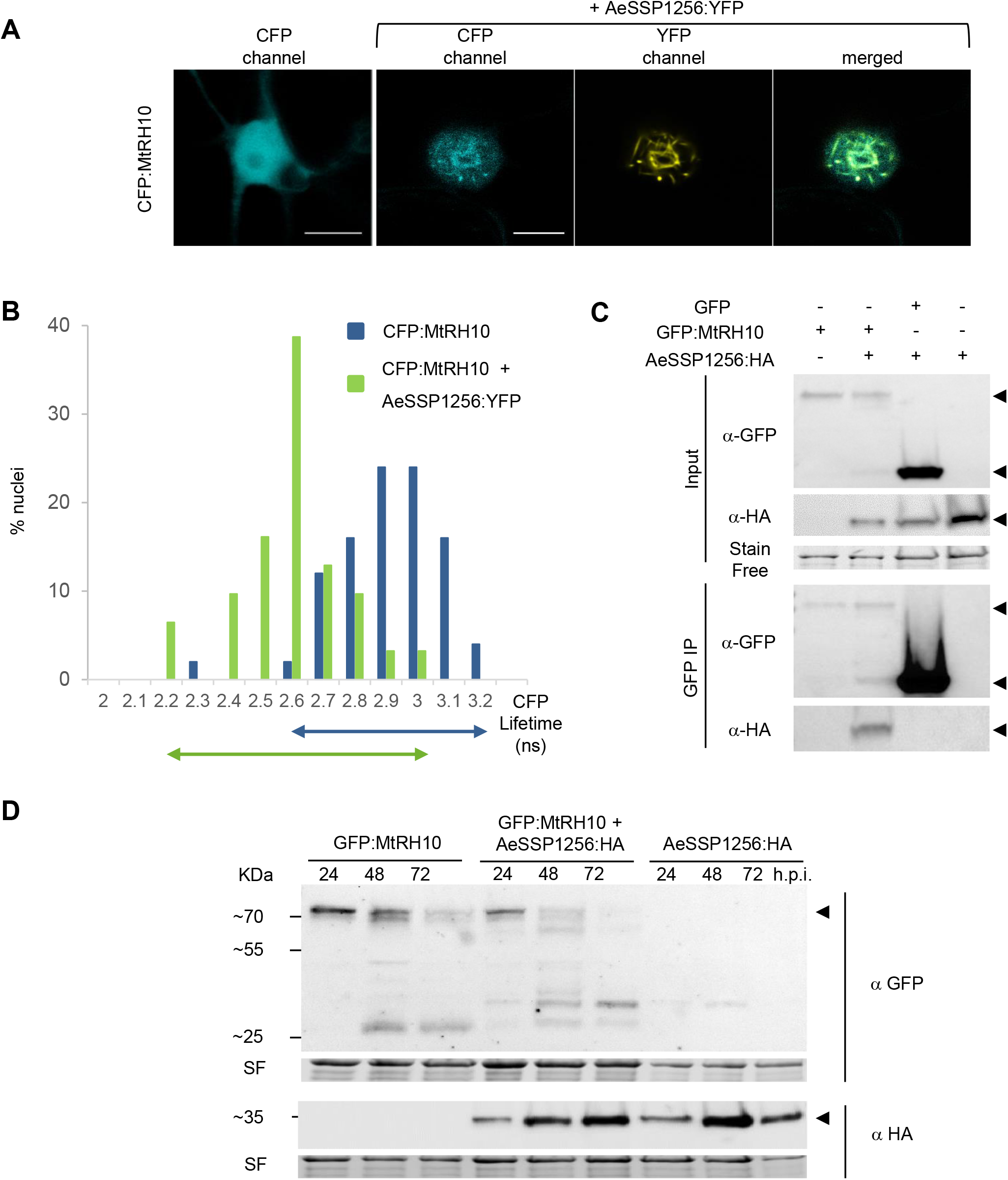
AeSSP1256 interacts and re-localizes the nuclear MtRH10 RNA Helicase around the nucleolus. **(A)** Confocal analyses on *N. benthamiana* agroinfiltrated leaves. The CFP:MtRH10 candidate presents a nucleocytoplasmic localization when expressed alone (Left panel), and is re-localized in the nucleus, mostly around nucleolus, in the presence of AeSSP1256:YFP proteins (Right panels). Pictures were taken at 24h post agroinfection. Scale bars: 10μm. **(B)** FRET-FLIM experiments indicate that CFP:MtRH10 and AeSSP1256:YFP proteins are in close association when co-expressed in *N. benthamiana* cells. Histograms show the distribution of nuclei (%) according to classes of CFP:MtRH10 lifetime in the absence (blue bars) or presence (green bars) of AeSSP1256:YFP. Arrows represent CFP lifetime distribution range. **(C)** Co-immunoprecipitation experiments confirm the direct association of the two proteins. Upper panel: anti-GFP and anti-HA blots confirm the presence of recombinant proteins in the input fractions. Lower panel: anti-GFP and anti-HA blots on output fractions after GFP immunoprecipitation. Arrows indicate the corresponding proteins. **(D)** anti-GFP and anti-HA blots on *N. benthamiana* leaf extracts expressing the GFP:MtRH10 alone or in combination with AeSSP1256:HA protein after 24, 48 or 72h post agroinfection. Arrows indicate the corresponding proteins. Note that GFP:MtRH10 is degraded faster in presence of AeSSP1256:HA.

To confirm this result, co-immunoprecipitation assays were carried out. A GFP:MtRH10 construct was co-transformed with AeSSP1256:HA construct in *N. benthamiana* leaves. As expected, the localization of GFP:MtRH10 protein in absence of AeSSP1256 was nucleocytoplasmic while it located around the nucleolus in the presence of the effector (**Supplemental Figure 4B**). Immunoblotting experiments using total proteins extracted from infiltrated leaves (24hpi) showed that AeSSP1256:HA proteins were co-immunoprecipitated with GFP:MtRH10, but not with the GFP alone (**Figure 4C**). These data indicate that AeSSP1256 associates with MtRH10 in the nucleus. To go further we checked the stability of the two proteins when expressed alone or in combination in *N. benthamiana* cells during 72 hours. While GFP:MtRH10 was still detected at 72h after agroinfiltration, it started to be degraded 48hpi (**Figure 4D**). Expression of the effector alone is stable along the time. In contrast, when the two proteins are co-expressed, GFP:MtRH10 is almost entirely processed at 48h, and no more detectable at 72h (**Figure 4D**), suggesting that the effector enhance instability of its host target. Taken together, these results strongly suggest an interaction between AeSSP1256 and two type of components, a ribosomal protein and a nuclear RNA helicase from *M. truncatula*.

### AeSSP1256 alters the RNA binding activity of MtRH10

DEAD-box RNA helicases are RNA binding proteins involved in various RNA-related processes including pre-rRNA maturation, translation, splicing, and ribosome assembly (Jarmoskaite and Russell, 2011). These processes are dependent to the RNA binding ability of the proteins. Therefore we checked whether MtRH10 is able to bind nucleic acids *in planta* using FRET-FLIM assays as described previously. As reported in **Table 3** and in **Figure 5A**, GFP lifetime of GFP:MtRH10 decreased in presence of the acceptor, from 2.32 ns +/- 0.02 to 2.08 ns +/- 0.03 due to FRET between GFP and Sytox, confirming as expected that MtRH10 protein is bounded to nucleic acids.

**Table 3:** FRET-FLIM measurements for GFP:MtRH10 with or without Sytox Orange, in presence or in absence of AeSSP1256:HA

| Donor | Acceptor | $\tau$ <sup>(a)</sup> | sem <sup>(b)</sup> | N <sup>(c)</sup> | E <sup>(d)</sup> | <sup>(e)</sup> p-value |
| --- | --- | --- | --- | --- | --- | --- |
| GFP:MtRH10 | - | 2.32 | 0.020 | 60 | - | - |
| GFP:MtRH10 | Sytox Orange | 2.08 | 0.027 | 60 | 10.3 | 1.30E<sup>-10</sup> |
| GFP:MtRH10<br>(relocalized) | -<br>(+ AeSSP1256:HA) | 2.30 | 0.023 | 60 | - | - |
| GFP:MtRH10<br>(relocalized) | Sytox Orange<br>(+ AeSSP1256:HA) | 2.30 | 0.020 | 60 | 0 | 0.789 |

$\tau$ : mean life-time in nanoseconds (ns). <sup>(b)</sup> s.e.m.: standard error of the mean. <sup>(c)</sup> N: total number of measured nuclei. <sup>(d)</sup> E: FRET efficiency in % : $E=1-(\tau_{DA}/\tau_D)$ . <sup>(e)</sup> p-value (Student's *t* test) of the difference between the donor lifetimes in the presence or absence of acceptor.

**Figure 5:**
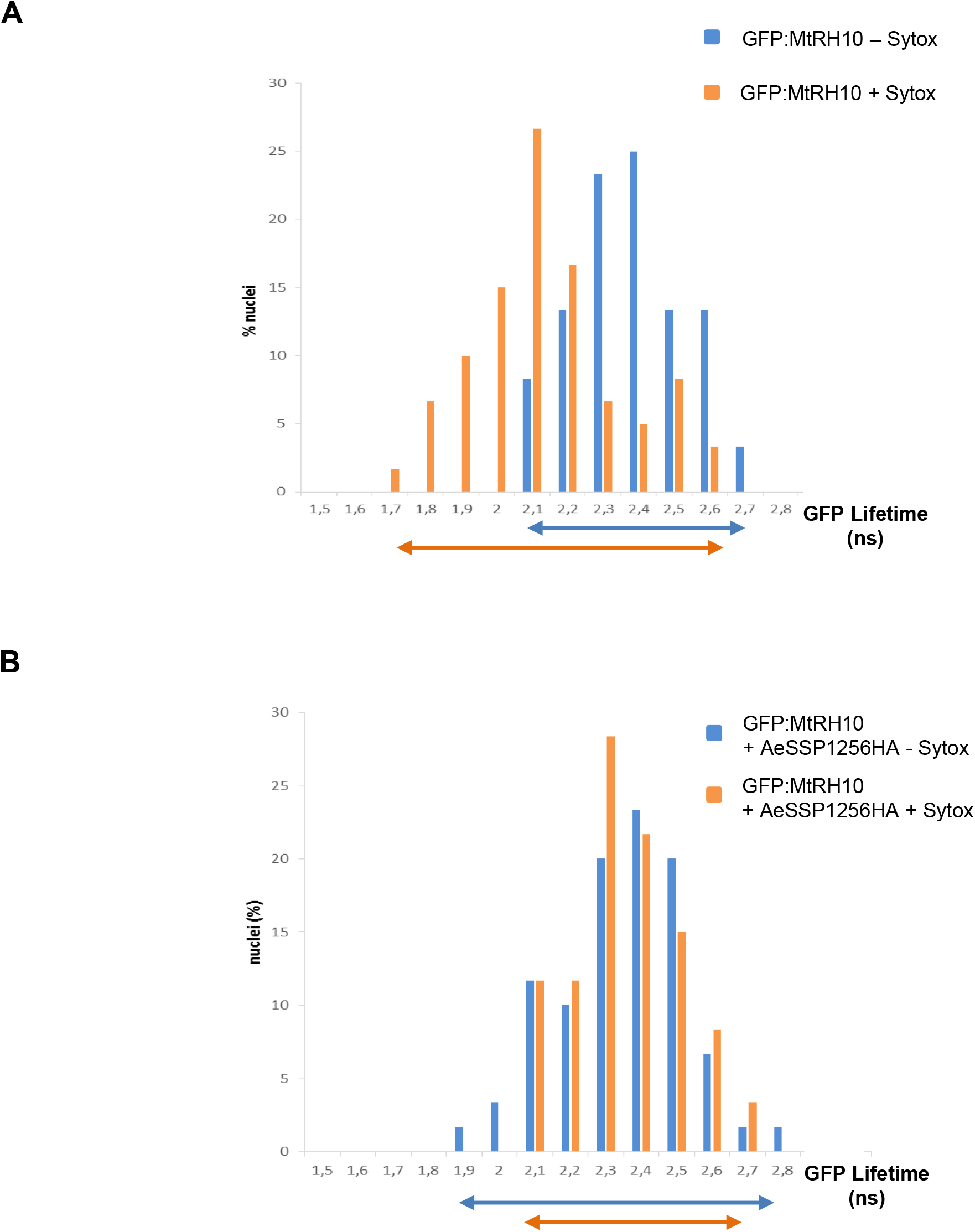
AeSSP1256 inhibits RNA binding activity of MtRH10. **(A)** FRET-FLIM experiments on *N. benthamiana* cells expressing GFP:MtRH10 in presence or absence of nucleic acids dye Sytox Orange. In presence of Sytox Orange, the GFP:MtRH10 lifetime decreases to shorter values, indicating that the proteins bounded to nucleic acids. **(B)** In presence of AeSSP1256:HA, when GFP:MtRH10 is re-localized around the nucleolus and interacts with AeSSP1256, no significant decrease in the GFP lifetime was observed in presence of Sytox Orange, meaning that the relocalized GFP:MtRH10 proteins were not able to interact with nucleic acids. Histograms show the distribution of nuclei (%) according to classes of GFP:MtRH10 lifetime in the absence (blue bars) or presence (orange bars) of the nucleic acids dye Sytox Orange. Arrows represent GFP lifetime distribution range.

To evaluate the role of AeSSP1256 on the function of MtRH10 we reasoned that the effector may perturb its binding capacity since it is required for the activity of numerous RH protein family (Jankowsky, 2011). We then co-expressed the GFP:MtRH10 construct with AeSSP1256:HA in *N. benthamiana* leaves and performed FRET-FLIM assays. Measurements made in nuclei where both proteins are detected due to the re-localization of MtRH10 indicated that GFP lifetime of GFP:MtRH10 remained unchanged with or without Sytox (2.3 ns in both conditions) showing that MtRH10 was not able to bind nucleic acids in the presence of the effector (**Table 3 and Figure 5B**). These data reveal that AeSSP1256 hijacks MtRH10 binding to RNA, probably by interacting with MtRH10.

### MtRH10 is expressed in meristematic root cells and its deregulation in *M. truncatula* impacts root architecture and susceptibility to *A. euteiches* infection

To characterize the function of MtRH10, we firstly consider the expression of the gene by mining public transcriptomic databases including Legoo (https://lipm-browsers.toulouse.inra.fr/k/legoo/), Phytozome (https://phytozome.jgi.doe.gov/pz/portal.html) and MedicagoEFP browser on Bar Toronto (http://bar.utoronto.ca/efpmedicago/cgi-bin/efpWeb.cgi). No variability was detected among the conditions tested in the databases and we do not detect modification of MtRH10 expression upon *A. euteiches* inoculation in our RNAseq data. To go further in the expression of the MtRH10 gene, transgenic roots expressing an MtRH10 promoter-driven GUS (β-glucuronidase) chimeric gene were generated. GUS activity was mainly detectable in meristematic cells, at the root tip or in lateral emerging roots (**Figure 6A**) suggesting a role in meristematic cell division. We complete MtRH10 analyses by overexpressing a GFP-tagged version in *Medicago* roots. The observation by confocal analyses of the subcellular localization of MtRH10 confirms its nucleocytoplasmic localisation as previously observed in *N. benthamiana* cells (**Figure 6B**). We also noticed the presence of brighter dots in the nucleolus corresponding probably to the fibrillar center. No developmental defects were detected in roots overexpressing MtRH10 (**Figure 6C-D**). To assess the effect of MtRH10 on root physiology and resistance to *A. euteiches*, a pK7GWiWG2:RNAi MtRH10 vector was design to specifically silence the gene in *Medicago* roots. RNA helicase gene expression was evaluated by qPCR 21 days after transformation. Analyses confirmed a reduced expression (from 3 to 5 times) compared to roots transformed with a GFP control vector (**Supplemental Figure 5**). Missense MtRH10 plants display a reduced number of roots coupled with shorter primary roots (**Figure 6C-D**) and a delay in development which starts with a shorter root apical meristem (RAM) (**Figure 6E-F**). This reduction in not due to smaller RAM cortical cell size (**Figure 6F**) suggesting a decrease in cell number. Longitudinal sections of roots expressing either RNAi MtRH10 or AeSSP1256 performed in elongation/differentiation zone (EDZ) revealed comparative defects in cortical cell shape or cell size (**Supplemental Figure 6A**). Cell area in missense MtRH10 or in AeSSP1256 roots is approximately reduced 2 times compared to GFP control roots (**Supplemental Figure 6B**) but proportionally the perimeter of those cells is longer than GFP cells, indicating a difference in cell shape (**Supplemental Figure 6C**). We noticed that most of EDZ cells in GFP roots present a rectangular shape, which seem impaired in missense MtRH10 and AeSSP1256 expressing roots. Thus we measured the perimeter-bounding rectangle (PBR) which calculates the smallest rectangle possible to draw with a given cell. A perimeter/PBR ratio of 1 indicates that the cell is rectangular. As presented in **Supplemental Figure 6D**, the perimeter/PBR ratio in GFP roots is close to 1 and significantly different than those observed in RNAi MtRH10 and AeSSP1256 roots. This analysis reveals that the reduction of MtRH10 expression or the expression of the effector AeSSP1256 in *Medicago* roots, impair the cortical cell shape. The similar phenotypic changes observed on MtRH10-silenced roots and AeSSP1256-expressing roots, suggests that the effector may affect MtRH10 activity in cell division regions of the roots.

**Figure 6:**
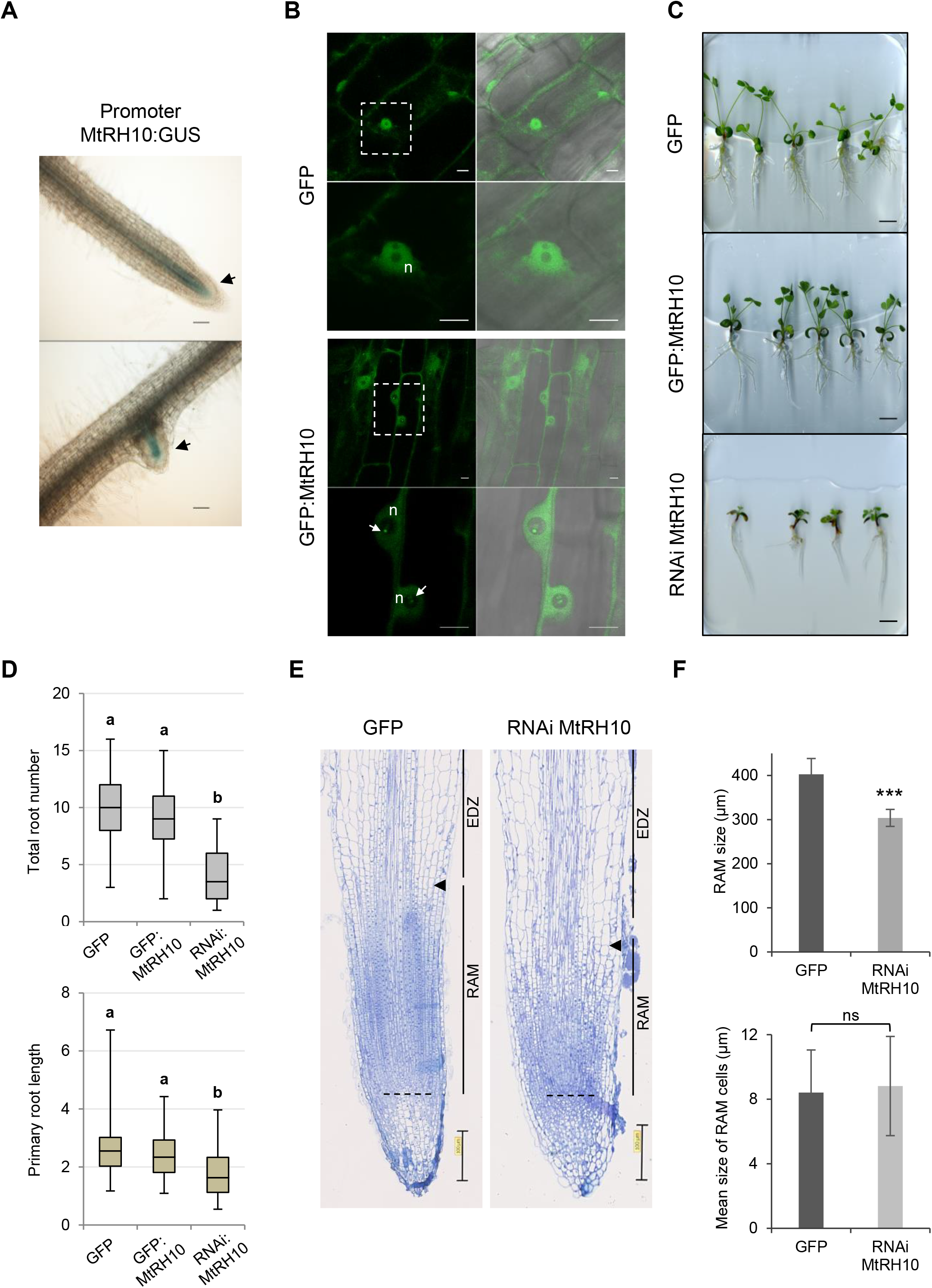
MtRH10 is expressed in meristematic cells of *Medicago truncatula* and its deregulation impacts root architecture. **(A)** GUS staining of MtRH10 promoter:GUS plants 21 d.a.t. Top panel: Root tip, bottom panel: emerging lateral root. Arrows indicate blue cells. Scale bars: 100μm. **(B)** Confocal pictures of *M. truncatula* roots transformed with GFP (top) or GFP:MtRH10 construct to overexpress MtRH10 (bottom). GFP:MtRH10 proteins harbor a nucleocytoplasmic localization with some brighter dots in the nucleolus (arrows). Lower panels represent nucleus enlargements. n: nucleus. Scale bars: 10μm. Left panel : 488nm, right panel: overlay (488nm + bright field). **(C)** Representative pictures of *M. truncatula* plants expressing either a GFP, a GFP:MtRH10 or RNAi MtRH10 construct 21 d.a.t. No particular phenotype was observed in the overexpressing MtRH10 plants. At the opposite, developmental delay appeared in missense MtRH10 plants. Scale bar: 1cm. **(D)** Total root number per plant (top) and primary root length per plant (bottom) in centimeters for GFP, GFP:MtRH10 and RNAi MtRH10 roots. Letters a and b indicate Student’s t-test classes (different classes if P < 0,01). **(E)** Representative longitudinal section of *M. truncatula* root tips expressing GFP or RNAi MtRH10 construct. Root apical meristem (RAM) size is determined from quiescent center (dot line) till the elongation/differentiation zone (EDZ), defined by the first elongated cortex cell of second cortical layer (arrowhead). Scale bars: 100μm. **(F)** Histograms of total RAM size and mean RAM cortical cell size. RAM of RNAi MtRH10 roots are smaller than in GFP control, but average cell size of cortical cells in RAM is not significantly different. Bars represent mean values and error bars are standard deviation. Asterisks indicate a significant p-value (t-test P < 0,0001, ns: not significant).

Having shown that MtRH10 is implicated in *M. truncatula* roots development, we test whether this biological function is related to pathogen colonisation. We therefore investigate by qPCR the presence of *A. euteiches* in silenced and overexpressed MtRH10 roots infected by the pathogen. As shown on **Figure 7**, overexpression of MtRH10 reduce the amount of mycelium in roots after 7, 14 and 21 dpi (1.8, 3.3 and 1.6 times less, respectively). We note by western-blot analyses a slight decrease in MtRH10 amount upon the time probably due to the accumulation of the AeSSP1256 effector (**Supplemental Figure 7**). As expected in roots where MtRH10 is silenced to 2 to 3 times as compared to GFP control roots, qPCR analyses revealed approximately 5 to 10 times more of the pathogen at 7, 14 and 21 dpi (**Figure 7**). Taken together these infection assays show that MtRH10 is involved in conferring basal resistance to *A. euteiches* at the root level.

**Figure 7:**
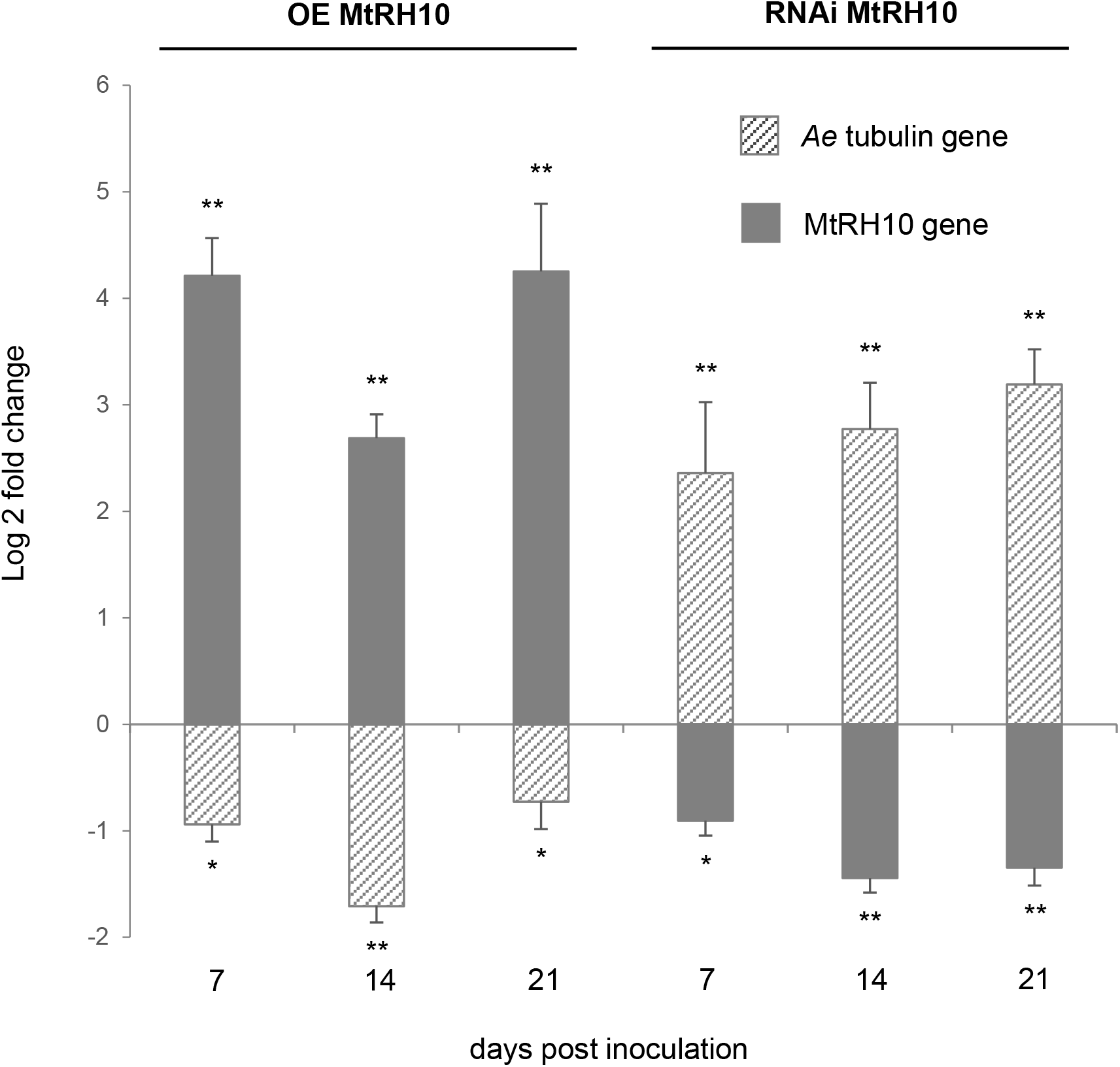
Deregulation of MtRH10 helicase gene expression in *Medicago truncatula* impacts *Aphanomyces euteiches* susceptibility. Expression values (Log2 fold change) for *A. euteiches* tubulin or MtRH10 genes in *M. truncatula* infected plants at 7, 14 and 21 d.p.i. in overexpressing GFP:MtRH10 plants (OE MtRH10) or in RNAi MtRH10 expressing plants compared to GFP control plants. Plants overexpressing MtRH10 gene are less susceptible to *A. euteiches* infection. In contrast, reduced expression of MtRH10 by RNAi enhances plant susceptibility to *A. euteiches*. Asterisks indicate significant differences (Student’s t-test; *: P < 0,05, **: p < 0,01). Bars and error bars represent respectively means and standard errors from three independent experiments. In total, N: 91 plants for GFP, 50 plants for GFP:MtRH10 and 50 plants for RNAi MtRH10 construct.

## Discussion

Protein effectors from filamentous plant pathogens such as fungi and oomycetes facilitate host colonization by targeting host components. However the molecular mechanisms that enhance plant susceptibility to the pathogen are still poorly understood. Here we report that the *A. euteiches* AeSSP1256 RNA-binding effector facilitate host infection by downregulating expression of plant ribosome-related genes and by hijacking from its nucleic target MtRH10, a *Medicago* nuclear RNA-helicase (RH). Thus the current study unravels a new strategy in which pathogenic oomycete triggers plant nucleolar stress to promote infection.

AeSSP1256 is an effector from the oomycete root pathogen *A. euteiches* previously shown to enhance oomycete infection (Gaulin et al., 2018). Despite the absence of any functional domain, in silico RGG/RG RNA-binding motif prediction (see for review (Thandapani et al., 2013)) prompt us to show by FRET/FLIM analysis that the secreted AeSSP1256 effector is an RNA-binding protein (RBP). RNAs play essential role in cell physiology and it is not surprising that filamentous plant pathogens may rely on RNA-dependent process to control host infection (for review see (Göhre et al., 2013; Pedersen et al., 2012)). Moreover RBPs are key players in the regulation of the post-transcriptional processing and transport of RNA molecules (Yang et al., 2018). However to our knowledge only three examples of RBPs acting as virulence factor of plant pathogens are known. This includes the glycine-rich protein MoGrp1 from the rice pathogen *Magnaporthe oryzae* (Gao et al., 2019), the UmRrm75 of *Ustilago maydis* (Rodríguez-Kessler et al., 2012) and the secreted ribonuclease effector CSEP0064/BEC1054 of the fungal pathogen *Blumeria graminis* which probably interferes with degradation of host ribosomal RNA (Pennington et al., 2019). This situation is probably due to the absence of conventional RNA-binding domain which render this type of RBP undetectable by prediction algorithms. The future studies that will aim to unravel the atlas of RNA-binding effector in phytopathogens should not only rely on computational analysis but will have to use functional approaches such as crystallization of the protein to validate function as performed with CSEP0064/BEC1054 effector (Pennington et al., 2019) screening method like the RNA interactome capture (RIC) assay develops in mammals (Castello et al., 2012) or FRET/FLIM assays to detect protein / nucleic acid interactions (Camborde et al., 2017).

We observed that when expressed inside roots of the partially resistant Jemalong A17 *M. truncatula* line, AeSSP1256 triggers developmental defects such as shorter primary roots and delay in root development. Defects in roots development and retarded growth are typical characteristics of auxin-related and ribosomal proteins mutants reported in *Arabidopsis* (Ohbayashi et al., 2017; Wieckowski and Schiefelbein, 2012). In addition, those composite *Medicago* promote infection of *A. euteiches*. This modification in the output of the infection is highly relevant since we previously observed that *M. truncatula* quantitative resistance to *A. euteiches* is correlated to the development of secondary roots (Rey et al., 2016). This activity is dependent on the nucleolar rim localization of AeSSP1256, closed to the nucleolus.

The nucleolus is a membrane-free subnuclear compartment essential for the highly complex process of ribosome biogenesis organized in three domains including the fibrillar center that contain rDNA, which are not yet engaged in transcription (reviewed in (Shaw and Brown, 2012). Ribosome biogenesis is linked to cell growth and required coordinated production of processed ribosomal RNA (rRNA), ribosomal biogenesis factors and ribosomal proteins (RP). In the nucleolus, ribosome biogenesis starts with the transcription of pre-rRNAs from rRNA genes, followed by their processing and assembly with RPs into two ribosome subunits (ie small and large subunit). In animals, perturbation of any steps of ribosome biogenesis in the nucleolus can cause a nucleolar stress or ribosomal stress which stimulates specific signaling pathway leading for example to arrest of cell growth (Pfister, 2019). The nucleolar rim localization of AeSSP1256 within the host cells suggested that this effector could interfere with ribosome biogenesis pathway to facilitate infection. This speculation was further strengthened by RNAseq experiments which showed that, within A17-roots, AeSSP1256 downregulated numerous genes implicated in ribosome biogenesis pathway, notably ribosomal protein genes. This effect was also detected in susceptible F83 *M. truncatula* lines infected by *A. euteiches* indicating that AeSSP1256, mimics some *A.euteiches* effects during roots invasion.

An Y2H approach led to the identification of putative AeSSP1256 plant targets and all but one correspond to predicted nuclear *M. truncatula* proteins. By a combination of multiple experiments as FRET-FLIM to detect protein/protein interactions, a L7 ribosomal protein (MtrunA17_Chr4g0002321) and a DExD/H box RNA helicase ATP-dependent (MtrunA17_Chr5g0429221) were confirmed as AeSSP1256-interacting proteins. The DExD/H (where x can be any amino acid) box protein family include the largest family of RNA-helicase (RH). Rather than being processive RH, several DExD/H box proteins may act as ‘RNA chaperone’ promoting the formation of optimal RNA structures by unwinding locally the RNA (for review see (Fuller-Pace, 2006)). These proteins are of major interest due to their participation to all the aspects of RNA processes such as RNA export and translation, splicing but the most common function of these proteins is in ribosome biogenesis including assembly (Jarmoskaite and Russell, 2011). Specific function of RH is probably due to the presence of a variable C-terminal ‘DEAD’ domain in contrast to the well conserved N-terminal ‘helicase core’ domain (for review see (Fuller-Pace, 2006)). This structural organization was detected in the MtRH10. This *M. truncatula* protein corresponds to the ortholog of the nucleolar human DDX47 (Sekiguchi et al., 2006), the nuclear yeast RRP3 (O’Day, 1996) and the nucleolar *Arabidopsis* AtRH10 RNA-helicases, all involved in ribosome biogenesis (Liu and Imai 2018; Matsumura et al. 2016), and the nucleolar rice OsRH10 (TOGR1) involved in rRNA homeostasis (Wang et al. 2016).

Like its human ortholog DDX47 (Sekiguchi et al., 2006), MtRH10 possesses a bipartite nuclear transport domain which can function as a nuclear localization signal (NLS) and two nuclear export signal (NES), and thereby it probably shuttles between the cytoplasm and the nucleus as reported for many others RNA helicases involved in rRNA biogenesis and splicing function (Sekiguchi et al. 2006; Wang et al. 2009). Fluorescence analysis showed a relocalization of the nucleocytoplasmic MtRH10 in the nucleoli periphery, when it is transiently co-express with AeSSP1256 in *N. benthamiana* cells. The change in MtRH10 distribution suggests that the interaction between the two proteins caused a mislocation of MtRH10 that can probably affect its activity. We thereby check the nucleic acid binding capacity of MtRH10 by FRET-FLIM approach. The decrease in the lifetime of GFP revealed the ability of MtRH10 to bind nucleic acids. Knowing that both proteins display the same properties, we further provided evidence that the presence of AeSSP1256 effector inhibits the nucleic binding capacity of MtRH10. This mechanism was also reported for the RNA-binding HopU1 effector from the plant bacterial pathogen *Pseudomonas syringae* which associates to the glycin-rich RNA binding 7 protein (GRP7) of *Arabidopsis* to abolish GRP7 binding to immune gene transcripts (ie FLS2 receptor, (Nicaise et al., 2013)). Here we cannot exclude that AeSSP1256 also blocks the putative helicase activity of MtRH10, but we favored an inhibitory mechanism of AeSSP1256 on MtRH10 activity as complex and at least in part due to both protein-protein interaction and nucleic acid interaction with the two proteins. Interestingly, we also noticed that co-expression of both proteins led to decrease in MtRH10 probably due to degradation of the protein. While this observation warrants further analyses, this effect is reminiscent of other effector activities which destabilize their targets (for review see (Langin et al., 2020)).

Plant genomes encode a large variety of DExD/H RH family in comparison to other organisms and numerous studies have shown that several are associated through their activity with plant development, hormone signaling or responses to abiotic stresses (for review see (Liu and Imai 2018)). Very few studies reported that DExD/H RH could also be involved in biotic stresses, like responses to pathogens. One example is the DExD/H RH OsBIRH1 from rice that enhanced disease resistance against *Alternaria brassicicola* and *Pseudomonas syringae* through activation of defense-related genes (Li et al. 2008). A recent study on oomycete reports the binding of the *Phytophthora sojae* RxLR PSR1 effector to a putative nuclear DExD/H RH. Although the affinity for nucleic acids was not evaluated for the RH, association of both partners promote pathogen infection by suppressing small RNA biogenesis of the plant (Qiao et al., 2015). Here we showed that MtRH10 knockdown tolerant A17 lines supported higher-level accumulation of *A. euteiches* in contrast to overexpressed MtRH10 lines, indicating the importance of MtRH10 for *M. truncatula* roots defense against soil-borne pathogens.

This works reveals that MtRH10 expression is restricted at the root apical meristematic zone (RAM) where cells divide (ie, primary and lateral roots). Missense MtRH10 roots harbor defects in the primary root growth and reduced number of roots. Longitudinal sections in elongation zone (EDZ) of these composite roots show a significant reduction in the size and shape modification of cortical cells indicating that MtRH10 is required for normal cell division. Defect in primary roots elongation is also detected in silenced AtRH10 and OsRH10 mutant (Matsumura et al. 2016; Wang et al. 2016). Thus MtRH10 plays a role on *Medicago* root development as its orthologs OsRH10 and AtRH10. At the cellular level we also observed in AeSSP1256-expressing roots, reduction in cell size in elongation zone, with defects in cell shape and in adhesion between cells of the cortex, maybe due to a modification of the middle lamella (Zamil and Geitmann, 2017). Thus AeSSP1256 triggers similar or enhanced effect on host roots development as the one detected in defective MtRH10 composite plants, supporting the concept that the activity of the effector on MtRH10 consequently leads to developmental roots defects. Several reports have indicated that *Arabidopsis* knockout of genes involved in rRNA biogenesis or in ribosome assembly cause abnormal plant development including restriction and retardation in roots growth (Ohtani et al., 2013; Huang et al., 2016, 2010). These common features suggest the existence of a common mechanism that regulate growth in response to insults of the ribosome biogenesis pathway, known as nucleolar stress response (for review see (Ohbayashi and Sugiyama, 2018)). How plant cells sense perturbed ribosome biogenesis and nucleolar problems is still an open question (Sáez-Vásquez and Delseny, 2019), but the ANAC082 transcription factor from *Arabidopsis* can be a ribosomal stress response mediator (Ohbayashi et al., 2017). In addition the recent report on the activity of the nucleolar OsRH10 (TOGR1, MtRH10 ortholog) implicated in plant primary metabolism through is activity on rRNA biogenesis, suggests that metabolites may play a role in this process. Finally our current study indicates that nuclear RNA-binding effector like AeSSP1256, by interacting with MtRH10, can act as a stimulus of the ribosomal stress response.

This work established a connection between the ribosome biogenesis pathway, a nuclear DExD/H RH, root development and resistance against oomycetes. Our data document that the RNA binding AeSSP1256 oomycete effector downregulated expression of ribosome-related genes of the host plant. The effector hijacked MtRH10, a nuclear DExD/H RH involved in root development, to promote host infection. This work not only provides insights into plant-root oomycete interactions but also reveals the requirement of fine-tuning of plant ribosome biogenesis pathways for infection success.

## Material and Methods

### Plant material, microbial strains, and growth conditions

*M. truncatula* A17 seeds were *in vitro*-cultured and transformed as previously described (Boisson-Dernier et al., 2001; Djébali et al., 2009). *A. euteiches* (ATCC 201684) zoospore inoculum were prepared as in (Badreddine et al., 2008). For root infections, each plant was inoculated with a total of 10μl of zoospores suspension at 10^5^ cells.ml^−1^. Plates were placed in growth chambers with a 16h/8h light/dark and 22/20°C temperature regime. *N. benthamiana* plants were grown from seeds in growth chambers at 70% of humidity with a 16h/8h light/dark and 24/20°C temperature regime. *E.coli* strains (DH5α, DB3.5), *A. tumefaciens* (GV3101::pMP90) and *A. rhizogenes* (ARQUA-1) strains were grown on LB medium with the appropriate antibiotics.

### Construction of plasmid vectors and *Agrobacterium*-mediated transformation

GFP control plasmid (pK7WGF2), +SPAeSSP1256:GFP and +SPAeSSP1256:YFP (named AeSSP1256:GFP and AeSSP1256:YFP in this study for convenience) and minus or plus signal peptide AeSSP1256:GFP:KDEL constructs were described in (Gaulin et al., 2018). Primers used in this study are listed in **Supplemental Table 3**. *M. truncatula* candidates sorted by Y2H assay (MtrunA17_Chr7g0275931, MtrunA17_Chr2g0330141, MtrunA17_Chr5g0407561, MtrunA17_Chr5g0429221, MtrunA17_Chr1g0154251, MtrunA17_Chr3g0107021, MtrunA17_Chr7g0221561, MtrunA17_Chr4g0002321) were amplified by Pfx Accuprime polymerase (Thermo Fisher; 12344024) and introduced in pENTR/ D-TOPO vector by means of TOPO cloning (Thermo Fisher; K240020) and then transferred to pK7WGF2, pK7FWG2 (http://gateway.psb.ugent.be/), pAM-PAT-35s::GTW:CFP or pAM-PAT-35s::CFP:GTW binary vectors.

Using pENTR/ D-TOPO:AeSSP1256, described in (Gaulin et al., 2018), AeSSP1256 was transferred by LR recombination in pAM-PAT-35s::GTW:3HA for co-immunoprecipitation and western blot experiments to create a AeSSP1256:HA construct and in pUBC-RFP-DEST (Grefen et al., 2010) to obtain a AeSSP1256:RFP construct for FRET-FLIM analysis. For RNAi of MtRH10 (MtrunA17_Chr5g0429221), a 328 nucleotides sequence in the 3’UTR was amplified by PCR (see **Supplemental Table 3**), introduced in pENTR/D-TOPO vector and LR cloned in pK7GWiWG2(II)-RedRoot binary vector (http://gateway.psb.ugent.be/) to obtain RNAi MtRH10 construct. This vector allows hairpin RNA expression and contains the red fluorescent marker DsRED under the constitutive *Arabidopsis* Ubiquitin10 promoter (http://gateway.psb.ugent.be/), to facilitate screening of transformed roots. For MtRH10 promoter expression analyses, a 1441nt region downstream of the start codon of MtRH10 gene was amplified by PCR (see **Supplemental Table 3**), fused to β-glucuronidase gene (using pICH75111 vector (Engler et al., 2014)) and inserted into pCambia2200:DsRED derivative plasmid (Fliegmann et al., 2013) by Golden Gate cloning to generate PromoterMtRH10:GUS vector.

Generation of *M. truncatula* composite plants was performed as described by (Boisson-Dernier et al., 2001) using ARQUA-1 *A. rhizogenes* strain. For leaf infiltration, GV3101 *A. tumefaciens* transformed strains were syringe-infiltrated as described by (Gaulin et al., 2002).

### Cross-section sample preparation for confocal microscopy

*M. truncatula* A17 plants expressing GFP or AeSSP1256:GFP constructs were inoculated with *A. euteiches* zoospores 21 days after transformation as indicated previously. Roots were harvested 21 days post inoculation, embedded in 5% low-melting point agarose and cutted using a vibratome (VT1000S; Leica, Rueil-Malmaison, France) as described in (Djébali et al., 2009). Cross-sections were stained using Wheat Germ Agglutin (WGA)-Alexa Fluor 555 conjugate (Thermo Fischer; W32464), diluted at 50 μg/ml in PBS for 30min to label *A. euteiches*.

### RNA-Seq experiments

Roots of composite *M. truncatula* A17 plants expressing GFP or AeSSP1256:GFP constructs were harvested one week later after first root emergence. Before harvest, roots were checked for GFP-fluorescence by live macroimaging (Axiozoom, Carl Zeiss Microscopy, Marly le Roi, France) and GFP-positive roots were excised from plants by scalpel and immediately frozen in liquid nitrogen. Four biological replicates per condition were performed (GFP vs AeSSP1256-expressing roots), for each biological replicate 20-40 transformed plants were used. Total RNA was extracted using E.Z.N.A.^®^ total RNA kit (Omega bio-tek) and then purified using Monarch^®^ RNA Cleanup Kit (NEB). cDNA library was produced using MultiScribe™ Reverse Transcriptase kit using mix of random and poly-T primers under standard conditions for RT-PCR program. Libraries preparation was processed in GeT-PlaGe genomic platform (https://get.genotoul.fr/en/; Toulouse, France) and sequenced using Illumina HiSeq3000 sequencer. The raw data was trimmed with trmigalore (version 0.6.5) (https://github.com/FelixKrueger/TrimGalore) with cutadapt and FastQC options, and mapped to *M. truncatula* cv. Jemalong A17 reference genome V. 5.0 (Pecrix et al., 2018) using Hisat2 (version 2.1.0) (Kim et al., 2019). Samtools (version 1.9) algorithms fixmate and markdup (Li et al. 2009) were used to clean alignments from duplicated sequences. Reads were counted by HTseq (version 0.9.1) (Anders et al., 2015) using reference GFF file. The count files were normalized and different expression were quantified using DESeq2 algorithm (Love et al., 2014), false-positive hits were filtered using HTS filter (Rau et al., 2013). GO enrichment were done using ErmineJ (Lee et al., 2005) and topGO (Alexa and Rahnenfuhrer 2020) software. RNASeq experiments on F83005.5 (F83) susceptible plants infected by *A. euteiches* and collected nine days after infection are described in (Gaulin et al., 2018).

### RNA extraction and qRT-PCR

RNA was extracted using the E.Z.N.A^®^ Plant RNA kit (Omega Bio-tek). For reverse transcription, 1μg of total RNA were used and reactions were performed with the High-Capacity cDNA Reverse Transcription Kit from Applied Biosystems and cDNAs obtained were diluted 10 fold. qPCR reactions were performed as described in (Ramirez-Garcés et al., 2016) and conducted on a QuantStudio 6 (Applied Biosystems) device using the following conditions: 10min at 95°C, followed by 40 cycles of 15s at 95°C and 1min at 60°C. All reactions were conducted in triplicates.

To evaluate *A. euteiches’s* infection level, expression of *Ae* α-tubulin coding gene (Ae_22AL7226, (Gaulin et al., 2008)) was analyzed and histone 3-like gene and EF1α gene of *M*. *truncatula* (Rey et al., 2013) were used to normalize plant abundance during infection. For *Aphanomyces* infection in plant over-expressing GFP, AeSSP1256:GFP or GFP:MtRH10, cDNAs from five biological samples were analyzed, given that a sample was a pool of 3 to 5 plants, for each time point, on three independent experiments, representing 45 to 75 transformed plants per construct. *M. truncatula* roots were harvested 7, 14 and 21 dpi. For missense MtRH10 experiments, downregulation of MtRH10 gene was first verified using cDNAs from five biological samples, given that a sample was a pool of 5 plants, harvested 21 days post transformation. For *A. euteiches* inoculation, three biological samples were analyzed, given that a sample was a pool of 3 plants, for each time point, on two independent experiments, representing around 50 transformed missense MtRH10 plants. Relative expression of *Ae* α-tubulin or MtRH10 helicase genes were calculated using the 2^−ΔΔCt^ method (Livak and Schmittgen, 2001). For qPCR validation of RNAseq experiment, cDNAs from five biological replicates (pool of three plants) of AeSSP1256-expressing roots were extracted 21 days post transformation. Primers used for qPCR are listed in **Supplemental Table 3**.

### Yeast Two Hybrid assays

An ULTImate Y2H™ was carried out by Hybrigenics-services (https://www.hybrigenics-services.com) using the native form of AeSSP1256 (20-208 aa) as bait against a library prepared from *M. truncatula* roots infected by *A. euteiches*. The library was prepared by Hybrigenics-services using a mixture of RNA isolated from uninfected *M. truncatula* F83005.5 (+/- 12%), *M. truncatula* infected with *A. euteiches* ATCC201684 harvested one day post infection (+/- 46%) and *M. truncatula* infected with *A. euteiches* harvested six days post infection (+/- 42%). This library is now available to others customers on Hybrigenics-services. For each interaction identified during the screen performed by Hybrigenics (65 millions interaction tested), a ‘Predicted Biological Score (PBS)’ was given which indicates the reliability of the identified interaction. The PBS ranges from A (very high confidence of the interaction) to F (experimentally proven technical artifacts). In this study we kept eight candidates with a PBS value from ‘A and C’ for validation.

### Analysis of amino acid sequence of MtRH10

Conserved motifs and domains of DEAD-box RNA helicase were found using ScanProsite tool on ExPASy web site (https://prosite.expasy.org/scanprosite/). MtRH10 putative NLS motif was predicted by cNLS Mapper with a cut-off score of 4.0 (Kosugi et al., 2009), and the putative NES motifs were predicted by NES Finder 0.2 (http://research.nki.nl/fornerodlab/NES-Finder.htm) and the NetNES 1.1 Server (la Cour et al., 2004).

### Immunoblot analysis

*N. benthamiana* leaves, infected *M. truncatula* roots or roots of *M. truncatula* composite plants were ground in GTEN buffer (10% glycerol, 25 mM Tris pH 7.5, 1 mM EDTA, 150 mM NaCl) with 0.2% NP-40, 10mM DTT and protease inhibitor cocktail 1X (Merck; 11697498001). Supernatants were separated by SDS-PAGE and blotted to nitrocellulose membranes. For GFP and GFP variant fusion proteins detection, anti-GFP from mouse IgG1κ (clones 7.1 and 13.1) (Merck; 11814460001) were used when monoclonal Anti-HA antibodies produced in mouse (Merck; H9658) were chosen to detect HA recombinant proteins. After incubation with anti-mouse secondary antibodies coupled to horseradish peroxidase (BioRad; 170-6516), blots were revealed using ECL Clarity kit (BioRad; 170-5060).

### Co-immunoprecipitation assay

Co-immunoprecipitation was performed on *N. benthamiana* infiltrated leaves expressing GFP, GFP:MtRH10 or AeSSP1256:HA tagged proteins. Total proteins were extracted with GTEN buffer and quantified by Bradford assay. 50μg of total proteins were incubated 3H at 4°C with 30μl of GFP-Trap Agarose beads (Chromotek; gta-20) under gentle agitation for GFP-tagged protein purification. After four washing steps with GTEN buffer containing 0,05% Tween-20, beads were boiled in SDS loading buffer.

### Confocal microscopy

Scanning was performed on a Leica TCS SP8 confocal microscope. For GFP and GFP variant recombinant proteins, excitation wavelengths were 488 nm (GFP) whereas 543nm were used for RFP variant proteins. Images were acquired with a 40x water immersion lens or a 20x water immersion lens and correspond to Z projections of scanned tissues. All confocal images were analyzed and processed using the Image J software.

### Cytological observations of transformed roots

Roots of composite plants expressing GFP, AeSSP1256:GFP, GFP:MtRH10 or RNAi MtRH10 were fixed, polymerized and cutted as described in (Ramirez-Garcés et al., 2016). NDPview2 software was used to observe longitudinal root sections of GFP or missense MtRH10 plants and to measure RAM size. Image J software was used for all others measurements. Average RAM cells size were estimated by measuring all the cells from a same layer from the quiescent center to the RAM boundary. Mean values were then calculated from more than 200 cells. In the elongation zone (EDZ) of GFP, AeSSP1256:GFP or missense MtRH10 roots, cell area and cell perimeter were measured in rectangular selection of approximately 300×600μm (two selections per root). To obtain a normalized cell perimeter, each cell perimeter is proportionally recalculated for a of 500μm^2^ area standard cell. To estimate cell shape differences, considering that cortical cells in EDZ of GFP control roots are mostly rectangular, we measured the perimeter bounding rectangle (PBR), which represent the smallest rectangle enclosing the cell. Then we calculated the ratio perimeter / PBR. Rectangular cells have a perimeter / PBR ratio close to 1. Three roots per construct from three independent experiments were used.

### FRET / FLIM measurements

For protein-protein interactions, *N. benthamiana* agroinfiltrated leaves were analysed as described in (Tasset et al., 2010). For protein-nucleic acid interactions, samples were treated as described in (Camborde et al., 2017; Escouboué et al., 2019). Briefly, 24 h agroinfiltrated leaf discs were fixed with a 4% (w/v) paraformaldehyde solution. After a permeabilization step of 10 min at 37°C using 200 μg/ml of proteinase K (Thermo Fisher; 25530049), nucleic acid staining was performed by vaccum-infiltrating a 5 μM of Sytox Orange (Thermo Fisher; S11368) solution. For RNase treatment, foliar discs were incubated 15 min at room temperature with 0.5 mg/ml of RNAse A (Merck; R6513) before nucleic acid staining. Then fluorescence lifetime measurements were performed in time domain using a streak camera as described in (Camborde et al., 2017). For each nucleus, fluorescence lifetime of the donor (GFP recombinant protein) was experimentally measured in the presence and absence of the acceptor (Sytox Orange). FRET efficiency (E) was calculated by comparing the lifetime of the donor in the presence (τ_DA_) or absence (τ_D_) of the acceptor: E=1-(τ_DA_) / (τ_D_). Statistical comparisons between control (donor) and assay (donor + acceptor) lifetime values were performed by Student *t-test*. For each experiment, nine leaf discs collected from three agroinfiltrated leaves were used.

### Accession Numbers

Transcriptomic data are available at the National Center for Biotechnology Information (NCBI), on Gene Expression Omnibus (GEO) under accession number [GEO:GSE109500] for RNAseq corresponding to *M. truncatula* roots (F83005.5 line) infected by *A. euteiches* (9dpi) and Sequence Read Archive (SRA) under accession number PRJNA631662 for RNASeq samples corresponding to *M. truncatula* roots (A17) expressing either a GFP construct or a native AeSSP1256:GFP construct. SRA data will be release upon acceptation of the manuscript.

## Supplemental Data

The following supplemental data are available:

**Supplemental Figure 1**: the nuclear localization of AeSSP1256 is required for biological activity in *M. truncatula* roots

**Supplemental Figure 2**: invasion of *M. truncatula* roots by the pathogen is unchanged in AeSSP1256 effector-expressing roots

**Supplemental Figure 3**: CFP:L7RP candidate and AeSSP1256:YFP are in close association

**Supplemental Figure 4**: AeSSP1256 drives the re-localisation of the nuclear MtRH10 RNA helicase, around the nucleolus in *N. benthamiana* cells

**Supplemental Figure 5**: Expression of MtRH10 is reduced in *M. truncatula* silenced-roots

**Supplemental Figure 6**: *M. truncatula* cell morphology is affected in RNAi MtRH10 and AeSSP1256:GFP expressing roots

**Supplemental Figure 7**: Western blot and confocal analyses on MtRH10-overexpressed roots infected by *A. euteiches*

**Supplemental Table 1**: RNASeq data of *M. truncatula* roots (A17) expressing either GFP construct or AeSSP1256:GFP construct. **ST1a**. Differentially expressed genes (DE), padj<0,0001. **ST1b**. Top10 GO of DE. **ST1c**. Venn diagram. **ST1d**. qRT-PCR.

**Supplemental Table 2**: Yeast two hybrid screening. **STE2a**. List of putative AeSSP1256 interactors after Y2H screening of *M. truncatula* roots infected by the pathogen. **ST2b**. FRET-FLIM validation of CFP:L7RP candidate

**Supplemental Table 3**: List of primers used in this study

## Contributions

LC designed, performed molecular approaches on AeSSP1256 and wrote the manuscript, AK prepared and analyzed the RNAseq-experiments performed in this study and wrote the manuscript, AJ and LC performed FRET/FLIM analyses, CP and LC developed confocal studies, ALR performed cross and longitudinal sections studies and analyzed roots architecture of the different samples, MJCP prepared and analyzed yeast two hybrid assay, performed candidates cloning. BD analyzed the data and wrote the manuscript. EG conceived, designed, and analyzed the experiments, managed the collaborative work, and wrote the manuscript. All authors read and approved the final manuscript.

## Acknowledgements

The authors would like to thanks the GeT-PlaGe genomic platform (https://get.genotoul.fr/en/; Toulouse, France) for RNASeq studies; H. San-Clemente and M. Aguilar for statistical analysis help (LRSV, France); S. Courbier and A. Camon for their assistance in cloning steps. This work was supported by the French Laboratory of Excellence project “TULIP” (ANR-10-LABX-41; ANR-11-IDEX-0002-02) and by the European Union’s Horizon 2020 Research and Innovation programme under grant agreement No 766048.

